# Quantifying similarities between MediaPipe and a known standard for tracking 2D hand trajectories

**DOI:** 10.1101/2023.11.21.568085

**Authors:** Vaidehi P. Wagh, Matthew W. Scott, Sarah N. Kraeutner

**Author notes:** Contributing authors.

## Abstract

Marker-less motion tracking methods have promise for use in a range of domains, including clinical settings where traditional marker-based systems for human pose estimation is not feasible. MediaPipe is an artificial intelligence-based system that offers a markerless, lightweight approach to motion capture, and encompasses MediaPipe Hands, for recognition of hand landmarks. However, the accuracy of MediaPipe for tracking fine upper limb movements involving the hand has not been explored. Here we aimed to evaluate 2-dimensional accuracy of MediaPipe against a known standard. Participants (N = 10) performed trials in blocks of a touchscreen-based shape-tracing task. Each trial was simultaneously captured by a video camera. Trajectories for each trial were extracted from the touchscreen and compared to those predicted by MediaPipe. Specifically, following re-sampling, normalization, and Procrustes transformations, root mean squared error (RMSE; primary outcome measure) was calculated for coordinates generated by MediaPipe vs. the touchscreen computer. Resultant mean RMSE was 0.28 +/-0.064 normalized px. Equivalence testing revealed that accuracy differed between MediaPipe and the touchscreen, but that the true difference was between 0-0.30 normalized px (t(114) = -3.02, *p* = 0.002). Overall, we quantify similarities between MediaPipe and a known standard for tracking fine upper limb movements, informing applications of MediaPipe in a domains such as clinical and research settings. Future work should address accuracy in 3-dimensions to further validate the use of MediaPipe in such domains.

## 1 Introduction

Within clinical and rehabilitation settings the evaluation and monitoring of a patient’s movement quality is important when assessing the effectiveness of a treatment [1]. Typically, this will involve standardized clinical assessments, which while helpful, often involve a degree of subjectivity (e.g., [2],[3]). For example, observing and rating movement quality based on a set criterion or scale, which may be susceptible to bias and inaccuracies [4]. Alternatively, motion capture systems can be used to provide more objective measurements of movement; however, challenges may arise due to cost and maintenance of equipment. Accordingly, a cost and time-effective way in which movement can be evaluated with high accuracy is desirable in clinical settings.

Marker-based motion tracking involves optical, mechanical, magnetic or inertial systems which need complex computational processing and resources [5]. Furthermore, the use of such marker-based systems can be troublesome when assessing populations with sensory difficulties (e.g., tactile sensitivity). Advances in computer vision technology have enabled ‘marker-less’ systems, eliminating the need for physical tracking equipment. Striking an optimal balance between the resources invested and the out-put gained is an important part of motion tracking software. The development of lightweight systems for motion tracking is one way of tailoring computational difficulty and accuracy according to its application. MediaPipe by Google https://developers.google.com/mediapipe is an open source pipeline framework for deploying customised machine learning and computer vision solutions for live and streamed data [6]. This pipeline has been used primarily for human pose estimation [7], object detection [8], image segmentation [9], with applications such as sign language recognition [10], land-mark detection for augmented reality solutions [11], interactive virtual reality solutions [12], and visually enabled robotic arm control [13] on embedded devices, websites and applications. Recent studies have focused on the use of MediaPipe in clinical research settings, with applications in rehabilitation [14], physiotherapy [15], posture recognition [16], human gait analysis [17], as well as diagnostics in telehealth [18]. Outcome parameters of research studies in these domains include trajectory analysis, range of motion analysis, task based assessment, and accuracy of classification models. While MediaPipe has been applied to gross upper limb movements for joint angle estimation [19], motor skill assessment [20], tremor identification [21], and biometric identification [22], it has not been optimized and validated for behavioural investigations of fine upper-limb movements. Validation of MediaPipe for this purpose would provide strong implications for cost-effective motion tracking in clinical settings, particularly for populations with fine upper-limb movement impairments (e.g, stroke survivors [23], individuals with Parkinson’s disease [24], cerebral palsy [25] or developmental coordination disorder [26]. Previous research has validated MediaPipe against known standards for full-body joint angle estimation with acceptable errors [27]. In addition, gross motor tracking accuracy using RGB-depth cameras and Mediapipe has been validated with low errors for lower limb movements in running [28] and stationary cycling [29], as well as in hip, knee, shoulder and elbow joint movements [30], but poor correlation with ground truth data is reported for ankle joint movements [29]. Hand skill motor assessment using similar depth sensing setups and MediaPipe yielded optimal results with errors lower than 1 cm [20], but disturbances in hand trajectories were reported in Chunduru et al. [31]. Fine upper-limb movements differ from these applications due to a large diversity in parameters relating to dexterity, speed, occlusions and overlaps that occur during movement and lower contrast patterns between individual features as compared to gross upper limb movements. Given the number of degrees of freedom, and complexity in assessing such fine movements, it is critical to determine and quantify the accuracy of MediaPipe in comparison with known standards. The current work aimed to evaluate whether the model solutions given by MediaPipe can be used for accurate tracking of 2-dimensional (2D), fine upper limb movements. To address this objective, we evaluated the 2D accuracy of MediaPipe against a known standard, using a touchscreen-based shape-tracing task. Specifically, this task was used to generate 2D trajectories of hand/arm movements on a touchscreen computer. Videos of the hand/arm movements were processed through the MediaPipe pipeline to obtain coordinates of the movement. Additional post-processing steps (re-sampling and normalization) were applied to standardize MediaPipe data by moving it into a common reference space. To assess the accuracy of MediaPipe predictions, the processed 2D coordinate data was compared to our known standard; coordinates obtained from the touchscreen computer. Following Procrustes transformations to facilitate comparison, root mean squared error (RMSE; our primary outcome measure) was obtained between MediaPipe and the touchscreen computer. We hypothesized that MediaPipe generated coordinates would be equivalent with those generated by the touchscreen computer.

## 2 Methods

### 2.1 Participants

Data were collected from 10 young healthy participants (aged 19.5±1.3, 9 females, 1 male, 9 right handed, 1 ambidextrous) who participated in the study. All participants had normal or corrected-to-normal vision, and were free of neurological disorders or any physical impairment that would impact upper-limb movement. Informed consent was obtained from all participants. Ethical approval was obtained from the Institution’s Research Ethics Board.

### 2.2 Behavioural Task

All participants engaged in a 2D shape-based tracing task performed on a touchscreen computer using custom software developed in Python programming environment (https://github.com/LBRF/TraceLab; Ingram et al. 2019). This task is described in Ingram et al. [32]. Briefly, each trial began with the participant tapping a ‘go’ button located on the lower corner of the screen to trigger movement of a white cursor originating from the starting point. Trajectories consisted of five-segments (curved paths between four additional points and ending at the starting point). All trajectories were animated clockwise from point to point, with no visual feedback (‘trace’) left on the screen for participants to track. Immediately following the animation, participants were instructed to reproduce the trajectory on the screen beginning and ending from the starting point, matching the speed at which it was animated. All participants performed the task in a seated position directly in front of a horizontally oriented 12.3” touchscreen (Microsoft Surface Pro 7) placed on a desk. Participants performed the task with the index finger of their dominant hand, with their non-dominant hand resting comfortably in their lap. All participants performed these tasks in blocks, comprising of 10 trials each. X-Y coordinates of the participant’s produced trajectory for each trial was recorded by the computer and stored for offline analysis. To obtain videos of the shape-tracing task, a camera (GoPro Hero8) was mounted directly above the touchscreen computer to simultaneously record performance of each trial. One video was recorded per each block (10 trials) and stored for offline analysis.

## 3 Data Analysis

### 3.1 Behavioural task analysis

Pre-processing of the touchscreen-based shape-tracing task data occurred as described in Ingram et al. (2019) [32]. Pre-processing was performed to standardize participant’s produced trajectories by optimally transforming response trajectories onto the original stimulus (animation) trajectories, to account for variability in timing (via dynamic time warping [33]; to allow for natural variation in movement speed dynamic time warping) and natural variations in movement (via Procrustes transformation [34]; to account for ‘shape accuracy’ independent of translation, rotation, and scale).

### 3.2 Video analysis

Videos were captured by a GoPro Hero8. Each video was cropped to 2-6 second clips (one trial per video). Twelve blocks of trials were chosen for final analysis, including one to two blocks for each participant. Five trials were excluded due to participant compliance, leaving a hundred and fifteen trials in our final analyses. OpenCV, an open-source computer vision and machine learning library [35] was used to extract frames from each 2-6 second clip in the form of ‘.jpg’ images. A mean of 174 frames were obtained per video depending on the duration of the participant’s tracing movement in each 2-6 second clip. These frames were evaluated using the MediaPipe Hands solution for Python.

### 3.3 MediaPipe Hands

MediaPipe Hands is a lightweight, customisable, high-fidelity hand and finger tracking solution that employs machine learning to detect 21 landmarks of a hand with right and left handedness detection. MediaPipe Hands uses a single shot palm detection model to detect hands in the input image, followed by a hand landmark detector model over the bounding box of the detected hand for precise key point location of 21 fingertip and knuckle coordinates in virtual coordinates and real-world coordinates. Twenty-one coordinates comprise of the fingertip and 3 knuckles of each finger, as well as a point centered at the base of the palm. The model is robust to self-occlusions, partially visible hands, and multiple instances of hands in a given image. For this experiment, the MediaPipe Hands legacy solution was deployed in a Google Colab environment (https://colab.research.google.com) to evaluate the input frames. The frames for each trial clip were iterated through and evaluated serially by the MediaPipe Hands model, and X-Y coordinates for dominant hand index finger tips were extracted frame-by-frame. This coordinate data was stored pertaining to each trial trajectory. The coordinates were captured between 0-1920 units of X-coordinates and 1080 units of Y coordinates for an image resolution of 1920 x 1080. X-Y coordinates of the index finger tip output for each trial were then post processed.

### 3.4 Post-processing

Our overall pipeline is shown in Figure 1. Output X-Y coordinates for all trials were 1) resampled to 1500 points using univariate spline interpolation with k = 3 (cubic spline interpolation, via the ‘Interpolated Univariate Spline’ function from the ‘scipy.interpolate’ library in Python; [36]), to interpolate between MediaPipe predicted points. These coordinates were obtained within the coordinate system bounded by the height and width of the input image, between axis limits of 1920x1080. 2) Resampled data was then normalised by minmax scaling in Python using the sk.preprocessing library [37] to obtain the final MediaPipe coordinate data between 0 and 1 for each trial. To facilitate accurate comparison of MediaPipe predictions with trajectories extracted from the touchscreen, coordinates generated from the touchscreen-based shape-tracing task were post-processed as described above. Specifically, trajectories extracted from the touchscreen-based shape-tracing task were also resampled to 1500 points, via the scipy.interpolate library, and normalized using the sk.preprocessing library given differences in reference frames and coordinate systems between Medi-aPipe and the touchscreen. Once data from both MediaPipe and the touchscreen-based shape-tracing task was extracted, resampled (1500 points) and normalised (range of 0 to 1), the two datasets were compared to quantitatively assess the accuracy of Medi-aPipe predictions using Procrustes transformations and analysis in R programming environment (version 4.2.1) [38] via the vegan package [39]. Specifically, geometric transformations are applied to the MediaPipe data set to rotate, scale, and translate the dataset to best match it to the touchscreen data set. The best match is determined by the lowest Root Mean Square Error (RMSE) between corresponding points of the two datasets. Figure 2 shows an exemplar trace with post-processing steps applied for illustrative purposes. Applying this Procrustes transformation and analysis, we obtained RMSE for each pair of coordinates predicted by MediaPipe and the data from our touchscreen computer (our known standard) for each trial.

**Fig. 1.**
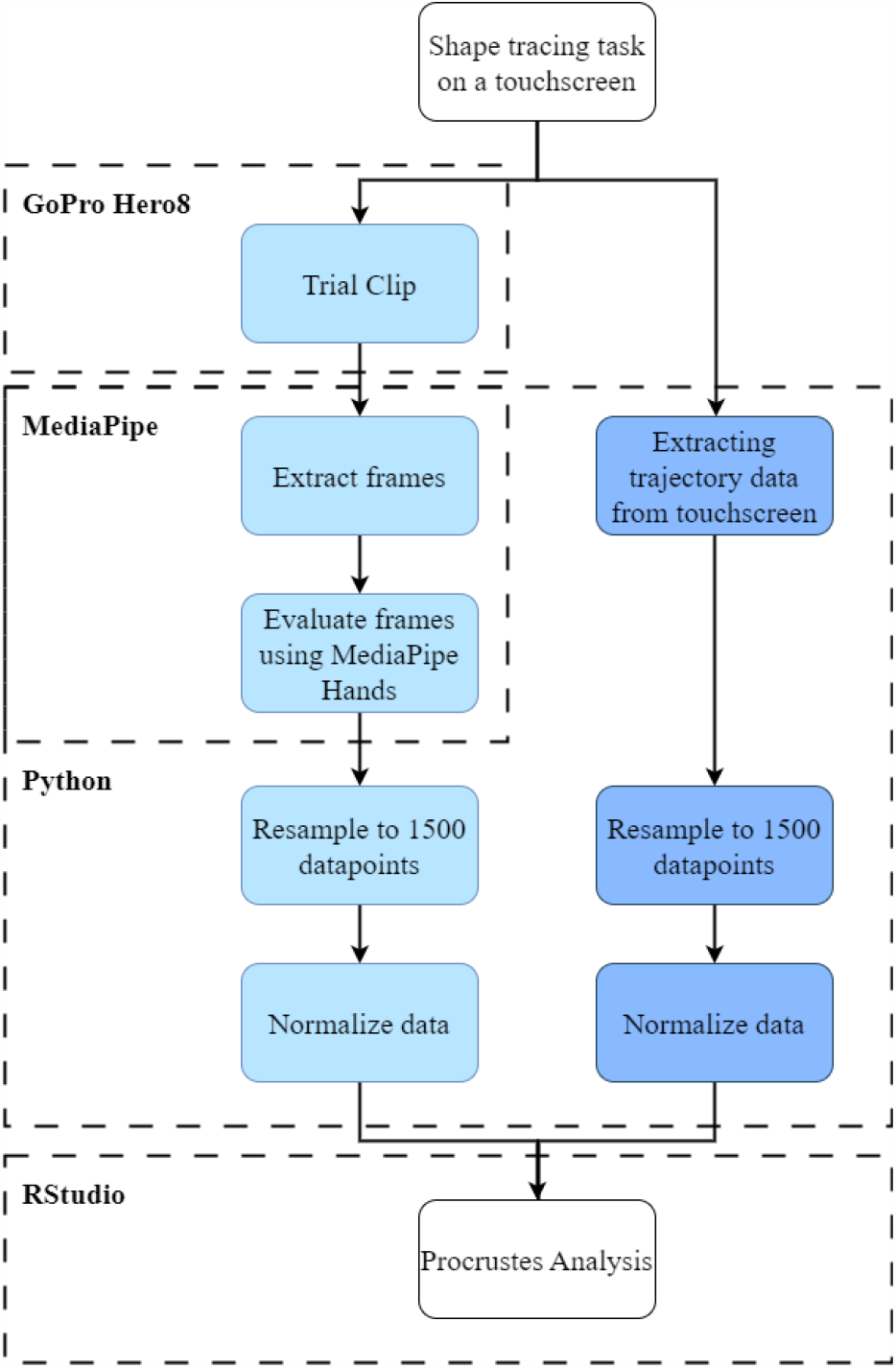
Pipeline Architecture

**Fig. 2.**
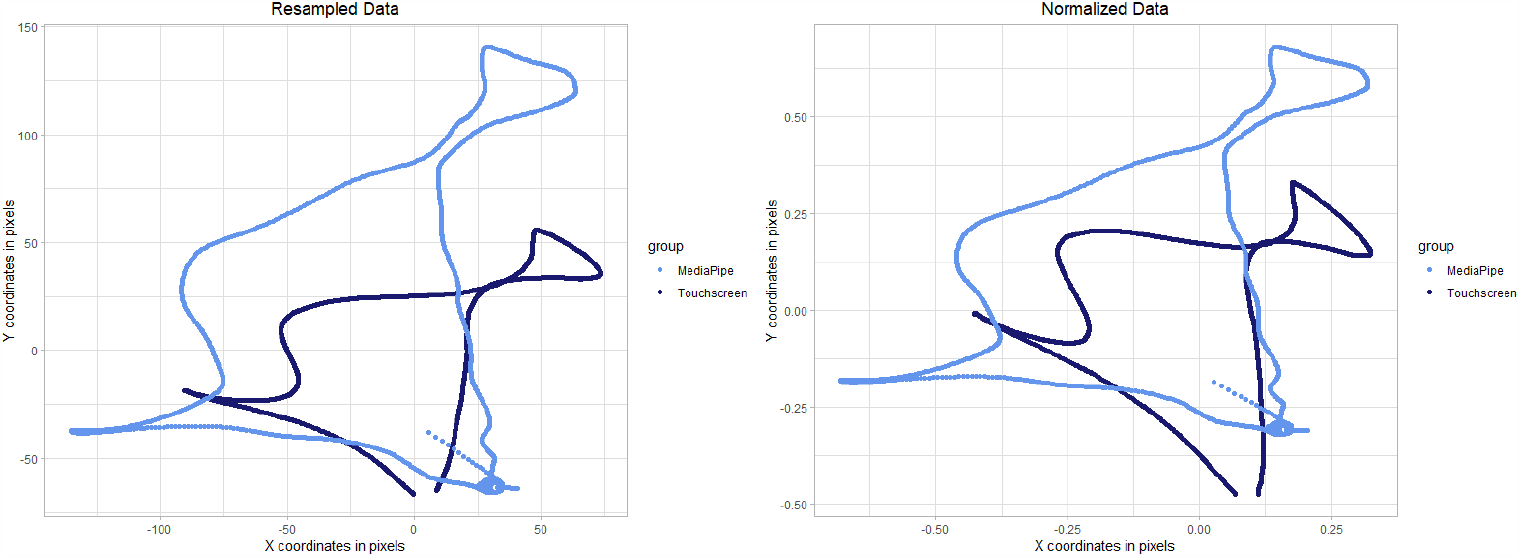
An exemplar trace predicted by MediaPipe (light blue) vs. data generated from the touchscreen-based shape-tracing task (dark blue). Traces are shown post Procrustes transformations after each post-processing step was applied on the data: re-sampling (left) and normalization (right).

### 3.5 Statistical analysis

#### 3.5.1 Procrustes RMSE

Equivalence tests were performed on the resultant RMSE calculated for coordinates generated by MediaPipe vs. the touchscreen computer, to determine whether MediaPipe coordinates were equivalent with touchscreen generated coordinates. Specifically, equivalence test was performed via two one-sided tests (TOST; *α <* 0.05) using the TOSTER package in R Programming Environment [40]. These tested the null hypotheses that true mean is equal to 0 (NHST), and true mean is more extreme than 0 and 0.30 (TOST).

#### 3.5.2 Exploratory analysis on impact of frame rates on RMSE

To assess the impact of the frame-rate parameter on the accuracy of tracing, we chose two blocks of trial videos each for three different frame rates - 30, 60, 120 FPS. We aimed to established a relationship between the frame rate of video capture and resultant Procrustes RMSE obtained for the blocks pertaining to each frame rate. We performed individual equivalence tests on this data and the results are tabulated in Table 1.

**Table 1.** Results of exploratory analysis.

| Frame rate | Mean RMSE | Std Deviation RMSE |
| --- | --- | --- |
| 30 | 0.29 | 0.042 |
| 60 | 0.28 | 0.056 |
| 120 | 0.32 | 0.043 |

Results of the equivalence tests conducted on Procrustes RMSE data (pixels) for trial blocks of data with 30, 60, and 120 frames-per-second (FPS)

## 4 Results

Visual inspection of our results on a frame-by-frame basis suggested that using Medi-aPipe to capture trajectories was overall similar to trajectories obtained from the touchscreen in short-duration clips, yet that post-processing resulted in greater overlap and improved the comparison. RMSE obtained by Primary Procrustes analysis was as follows: Mean RMSE = 0.28 +/- 0.064 px. The equivalence test was significant, t(114) = -3.02, *p* = 0.002 (mean = 0.28 90% C.I.[0.27, 0.29]; Hedges’s g = 4.40 90% C.I.[3.96, 5.0]). At the selected error rate, the true mean was found to be between 0 and 0.3. In the exploratory analysis, the lowest RMSE was observed at a frame rate of 60 FPS (Mean: 0.28 +/-0.056 px) for which the equivalence test was significant, t(19) = -1.89, p = 0.037 (Mean = 0.27 90% C.I.[0.26, 0.30]; Hedges’s g = 4.73 90% C.I.[3.70, 6.54]).

## 5 Discussion

As a proof of concept, we evaluated the extent to which MediaPipe-predicted trajectories deviated from those generated by a touchscreen computer (a “gold standard”). In partial support of our hypotheses, while a statistical difference between these trajectories was found, this difference was limited. These results support the accuracy of MediaPipe in determining hand poses and extracting coordinate data in complex poses with self occlusion. For example, accurate coordinates can be extracted when key features of hands are occluded, such as the palm by the fingers while making a fist. This also holds true in the presence of multiple hands and from low resolution or blurred images. However, it is important to consider the trade off between the accuracy and computational complexity of motion tracking systems to conclusively determine the application of MediaPipe in research and clinical domains such as physiotherapy, rehabilitation, and tele-rehabilitation. Eliminating the need for costly and complex physical tracking equipment enables motion tracking in a wide-variety of clinical domains where it may not always feasible nor possible to use a marker-based system. Validating such markerless systems is the first step towards bridging the gap between theory and practice of remote clinical assessment.

Our results address the research gap in fine limb tracking validation in 2D space, by providing a quantitative assessment of RMSE for complex tracing paths informing on future applications of MediaPipe in these domains. Our pipeline evaluated videos of 2-6 second duration with each video representing one trial, with a mean of 174 frames evaluated for each video. The reported error can be attributed to distortions during video capture, approximated geometric transformations or low resolution data capture. Our secondary, exploratory analysis discusses the optimal values of the frame-rate parameter to increase accuracy of the pipeline. We suggest the use of 60 frames-per-second to for recording the optimal videos. In our pipeline, a low frame rate of 30 FPS gave acceptable results, but a higher frame rate of 120 FPS gave increased errors associated with increased re-sampling points. An optimal frame rate value of 60 FPS is standard for most recording devices available today.

Our findings also suggest that post-processing steps should be included in an optimal MediaPipe pipeline for investigations of fine upper-limb movement. Here, post processing methods including re-sampling and normalisation improved transformation output, and in particular adjusted outputs to a common coordinate system to enable comparison with other systems (in this case, the touchscreen computer) or to provide a method of standardization across participants. Importantly, for videos captured with lower FPS, re-sampling can bridge the gap between low-cost data capture methods and highly accurate data. Normalisation can further help eliminate the need of shifting coordinate systems before comparison. Post processing data obtained from MediaPipe thus helps refine the output and bring about accurate output with low-cost motion capturing systems. With most modern cameras, 60 FPS is standard. Successful motion tracking has been presented for full body posture detection [16] and depth inclusive hand motor skill assessment [20] with 30 FPS. Higher FPS can also provide more data for key-point and pose extraction, leading to increased number of data points and thus, accuracy. Slow motion videos captured with 180 FPS and beyond as seen in [21] can also eliminate the need for re-sampling, but they come with high data capture costs. Such high FPS cannot be evaluated in real time, as standard webcams are not capable of capturing slow motion videos. Customizing the post processing pipelines according to a particular task is also crucial to the application of MediaPipe in different domains.

It is important to consider that our investigation comprised a typical population and was restricted to 2D coordinates. Nevertheless, these findings yield positive implications for outcome-based assessments of movement in typical and potentially clinical populations, particularly those with fine motor impairment (e.g., stroke survivors, individuals with cerebral palsy, developmental coordination disorder and those with Parkinson’s disease). MediaPipe provides a powerful model that is also capable of extracting 3D poses from 2D image inputs of hand poses. While showing 2D accuracy of MediaPipe3D represents an important first step towards its validation, future work is needed to assess its accuracy and feasibility for tracking 3D fine upper-limb movements, which when coupled with this 2D analysis, may provide a comprehensive assessment of movement form and outcome. Furthermore, equipment-free movement assessment tools such as MediaPipe provide a fruitful avenue for the assessment of home-based – upper limb focused – therapies which show to be effective (e.g., [41]; [42]; [43]; [44]). For instance, patients may record themselves performing a task via mobile phone and send these to their physiotherapist for documentation and assessment. Here, we tested MediaPipe as a marker-less motion capture system in a laboratory setting, informing on its use in clinical settings. MediaPipe also offers solutions in pose estimation for camera input, video input and livestream input for various mobile, web or device applications. Testing MediaPipe in more ecological settings is the next step, that may enable highly customisable motion tracking in variable environments in terms of backgrounds, obstructions and light illuminations. Robustness of the models to such variances is a crucial feature of motion tracking systems to ensure applicability in diverse settings.

## 6 Conclusion

In this study, we assessed the 2D accuracy of MediaPipe, a novel, marker less pose estimation tool, against a known standard. Our findings show an equivalence (within a range of 0 to 0.30 normalised px) between trajectories predicted by MediaPipe and those extracted from a touchscreen-based shape-tracing task. Our exploratory analysis suggests the use of a standard frame-rate of 60 FPS to replicate this methodology. Future work should address low cost motion capture using 2D image data for 3D pose estimation to further validate the use of MediaPipe in clinical populations and different settings (e.g., clinical, research and diagnostic settings). While, this work represents a necessary step towards validating the use of MediaPipe in investigations of fine upper-limb movement, future work is required to assess accuracy of MediaPipe for tracking 3D fine upper-limb movements across a larger sample.

## Declarations

### Ethics approval

Ethical approval was obtained from the Institution’s Research Ethics Board.

### Consent to participate

Informed consent was obtained from all participants.

### Consent for publication

All authors approved of the manuscript in its current form.

### Availability of data and materials

Data will be made available upon reasonable request to the corresponding author.

### Code availability

The code used to analyze these data is available at [insert github link here].

### Authors’ contributions

VW: data curation, methodology, software, formal analysis, visualization, writing – original draft, writing – review & editing.

MS: methodology, formal analysis, writing – review & editing.

SK: conceptualization, methodology, writing – original draft, writing – review & editing, funding acquisition, supervision, project administration.

